# Synaptic propagation in neuronal networks with finite-support space dependent coupling

**DOI:** 10.1101/2020.10.28.359877

**Authors:** Ricardo Erazo Toscano, Remus Osan

## Abstract

Traveling waves of electrical activity are ubiquitous in biological neuronal networks. Traveling waves in the brain are associated with sensory processing, phase coding, and sleep. The neuron and network parameters that determine traveling waves’ evolution are synaptic space constant, synaptic conductance, membrane time constant, and synaptic decay time constant. We used an abstract neuron model to investigate the propagation characteristics of traveling wave activity. We formulated a set of evolution equations based on the network connectivity parameters. We numerically investigated the stability of the traveling wave propagation with a series of perturbations with biological relevance.

## 2 Introduction

Traveling waves of electrical activity are everywhere in the nervous system. For example, cortical brain waves of activity are found in visual [1–6], olfactory [7, 8], auditory [9, 10], and motor [11, 12] cortices, as well as in the cerebellum [13]. Traveling waves may play roles in working memory [14], sensory processing, phase coding [15, 16], and sleep [17]. In the hippocampus, traveling waves of synchronization, such as theta and gamma waves, influence spatial-temporal dynamics [18, 19]. Sensorimotor systems rely on traveling waves up and down the spinal cord [20], and sensory systems such as in the invertebrate olfactory network [21], cuttlefish skin pigmentation [22]. Understanding the dynamics of traveling waves bridges a gap between observable phenomena and neuroscience theory.

In the present paper, we investigate traveling waves in simplified neuronal networks. We restrict the analysis of single-spike integrate-and-fire neuronal network activity propagation [23]. Our model assumes that synaptic connections between neurons are space-dependent: the finite support connectivity kernel assumes that synaptic coupling does not decay with distance. One neuron is coupled to all neighbor neurons within one synaptic space constant (synaptic footprint *σ*). We formulated a system of ordinary differential equations for traveling wave propagation to study the first, second, and potentially higher-order derivatives of firing times as a function of space. The finite support connectivity kernel and the time evolution of spike-dependent synaptic excitation are present in the evolution equations in an analytically tractable form. The evolution equations show that traveling wave dynamics depend on local dynamics and other parameters with a delayed effect. In essence, the excitation from one neuron affects all coupled neurons within one synaptic footprint *σ* and has delayed effects beyond its spatial boundaries. A key characteristic of our model is that the evolution equations are linear and can be solved analytically. The finite support connectivity function is mathematically and conceptually more straightforward than other connectivity functions widely used, such as Gaussian [24] or exponential decay [23]. However, constant speed wave dynamics in finite support connectivity kernel are more complicated than waves in networks with exponential decay kernel. Specifically, for the exponential kernel, wave acceleration depends quadratically on instantaneous speed plus delayed parameters, a system with straightforward dynamics [23]. The approach studied in the present paper allows us to solve equations for speed and acceleration explicitly and to define precisely how wave speed and acceleration fluctuate as a function of time and space. The derivations of our equations of evolution are in agreement with the numerical simulations. The evolution equations predict the traveling wave speed and acceleration based on the network excitability parameters. We further investigated the dynamics of traveling waves with different biologically relevant perturbations; these show that traveling waves are all-or-none events: waves can only propagate at the speed solution determined by the excitability parameters, or fail to propagate and die off.

## 3 Wave evolution in Integrate-and-Fire neuron model

Our theoretical framework accurately describes the system of activity propagation in neuronal networks; the framework has simple assumptions about neuronal excitability and network connectivity. In our simulations, we initiated traveling wave activities by applying an external current to a subset of neurons in the network labeled the “shocked region.” Consequently, the neurons in the shocked region spike at the same time (t=0). For simplicity, we assume the wave propagates only in the positive direction, and we monitor the spiking times of the neurons in the network. As neurons integrate excitatory synaptic inputs, they may reach the firing threshold and spike. Here the spiking times of neurons are a monotonic function of their position ‘x’ within the network. For numerical simulations, neurons in the network are located adjacent to one another and separated by a discretization constant *δ*. For a single spike-wave, we get a starting equation of the integral form

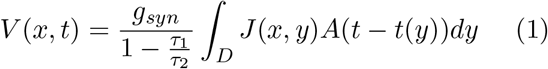

where

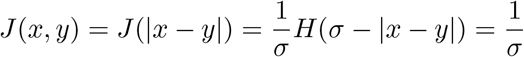

if |*x* − *y*| < *σ* and 0 otherwise

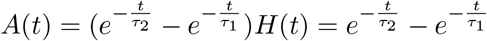

if *t* > 0 and 0 otherwise. With 0 < *τ*_1_ < *τ*_2_ and *σ* > 0

The consistency equation [Eq. 2] describes membrane potential (V) as a function of time and space-dependent kernel multiplied by network excitability parameters: 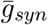, *σ*, *τ*_1_, *τ*_2_ (synaptic conductance, synaptic space constant, membrane time constant, and synaptic decay time constant). The neurons are set to spike when V = V_*T*_; the crossing of the threshold results in additional excitation. The parameters of the finite support kernel ‘x’ and ‘*x − σ*’ define the boundaries of the integral. [Eq. 2]:

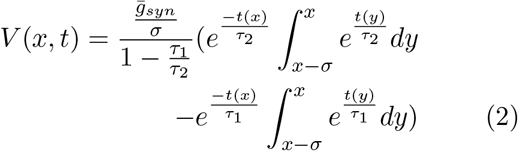

The wave instant velocity (equation 3) is the inverse of the first derivative of equation 2 with respect to ‘x’:

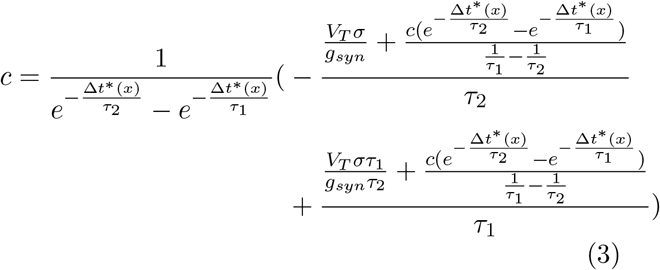

where we use the notation

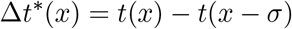

Figure 1 shows that the wave speed observed in numerical simulations and the solution of equation 3 show remarkable agreement. The wave speed evolves toward c_*fast*_ and oscillates transiently.

**Figure 1.**
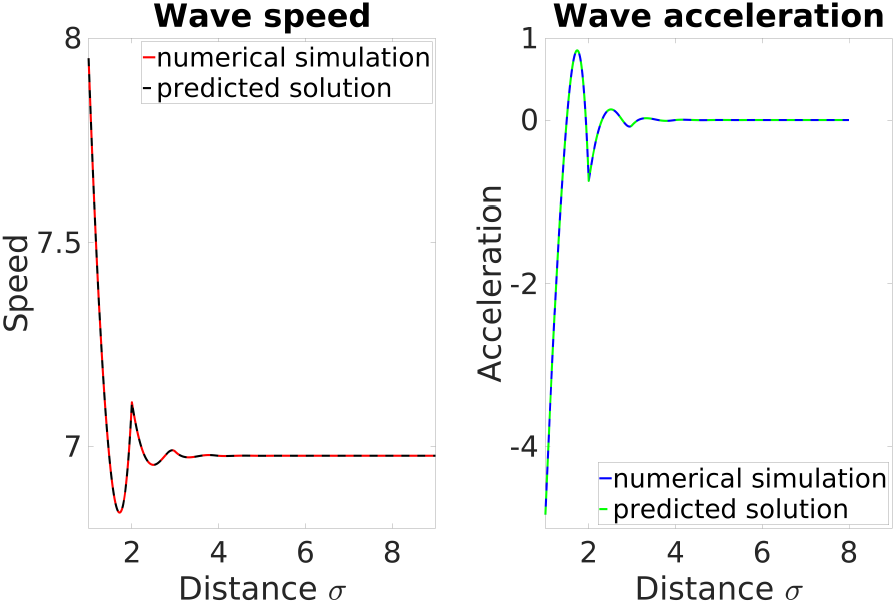
Comparison of analytical solutions and numerical simulations: The finite support neuronal network of integrate-and-fire neurons with excitatory coupling is at rest; at t=0, the neurons at x=0 to x=1 receive an additional current that drives them over the threshold. All shocked neurons spike synchronous because they receive their input at the same time. The wave evolution shows damped oscillations with an amplitude that decays exponentially. The self-propagating wave settles at the constant speed c_*fast*_. The left panel shows the wave speed as a function of space. The red trace is the computer simulation, and the black trace is the speed solution from equation 3. The right panel shows traveling wave acceleration as a function of space. The blue trace is the computer simulation, and the green trace is the solution from equation 4. Parameters: 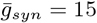, *σ* = 1, *τ*_1_ = 1, *τ*_2_ = 2, *V_T_* = 1.

We can determine 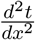 as a function of speed and convert 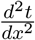 into the instantaneous acceleration. We used the same relationship from Zhang and Osan [56]:

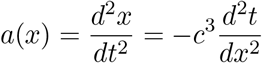

The wave instantaneous acceleration is:

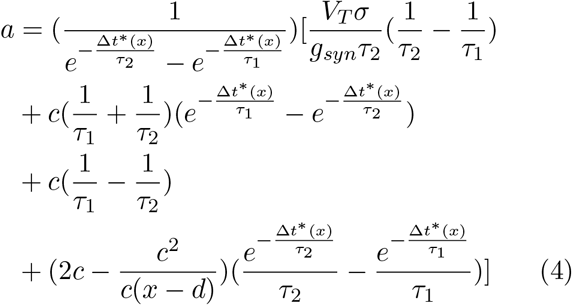

The equations of evolution describe traveling wave propagation in neuronal networks with finite support connectivity. The system of equations has the following unknown variables: equation 3 has t(x) and t(x-*σ*), the spike time of neuron at the location ‘x’ and spike time of neuron at the location ‘x-*σ*’; equation 4 has c(x) and c(x-*σ*), the speed of the wave at the location ‘x’ and the speed at the location ‘x-*σ*.’ We further explored the equations of evolution to estimate the speed of the traveling wave. We assume two intuitive features: (1) self-propagating traveling waves move at a constant speed, and (2) traveling waves that do not self-propagate eventually extinguish. The acceleration of constant speed waves is zero; therefore solving equation 4 when a=0 yields the consistency equation (equation 5), which describes neuron membrane potential as a function of traveling wave speed:

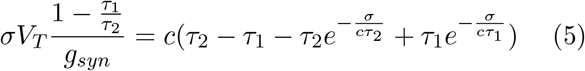

For a traveling wave that arrives from −∞ to a location ‘x’, and with speed c, one can compute the corresponding voltage. If the voltage is *V_T_* then the solution c is consistent with Eq. 5; otherwise, a traveling wave with speed c cannot exist.

The accuracy of the equations of evolution allows us to estimate the solutions of traveling wave propagation. Figure 2 illustrates that *c*_*fast*_ and c_*slow*_ arise in the intersection between V_*T*_ and V(c). Figure 2 also shows how the wave acceleration and speed oscillate while the wave evolves towards the stable state of activity propagation, where a=0 and c=c_*fast*_.

**Figure 2.**
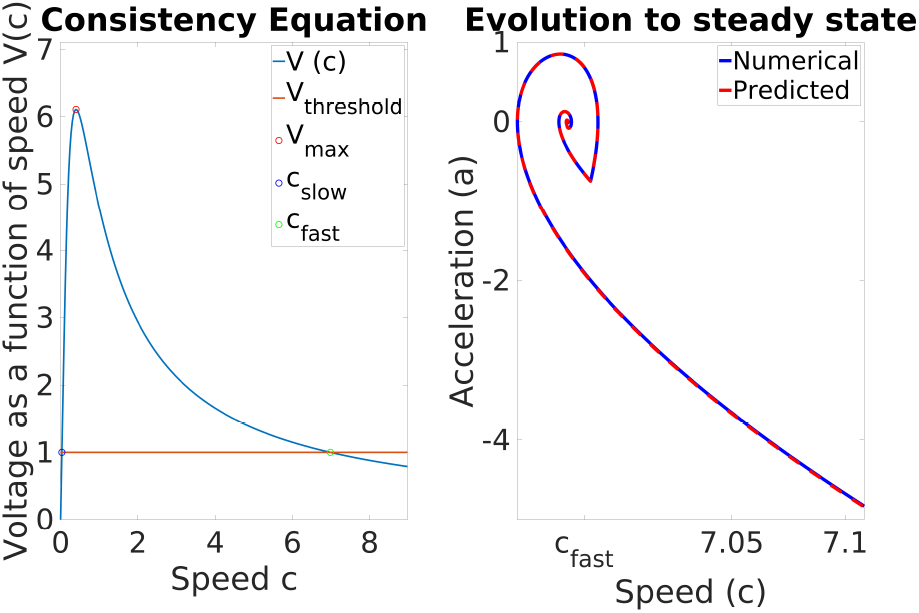
Stable and unstable fixed points of the wave propagation system. The left panel graphs the solutions to the consistency equation as intersections between V = V(c) and the horizontal line at *V_T_*. According to the consistency equation, the neuron membrane potential is a function of traveling wave speed. These numerical solutions allow one to compute both c_*fast*_ and c_*slow*_. *V_max_* denotes the maximum achievable voltage as a function of the traveling wave speed. If this value is less that *V_T_* no traveling wave can exist. The right panel shows results from numerical simulations. The attractor graph of acceleration as a function of speed indicates that the traveling wave settles into a constant speed solution. The magnitude of the oscillations is consistent with the synaptic footprint space constant *σ*. The speed (c) and acceleration (a) of the traveling wave oscillate while approaching the intersection between c= c_*fast*_, and a=0. Parameters: 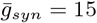, *σ* = 1, *τ*_1_ = 1, *τ*_2_ = 2, *V_T_* = 1.

## 4 Perturbation-based stability analysis

First, we established the existence of constant speed traveling waves in neuronal networks of Integrate-and-Fire neurons with finite support connectivity kernel. Next, we use a series of approaches to investigate the stability of the constant speed traveling wave. Here, the stable speed solution (c_*fast*_) is computed from equation 5 as well as determined from numerical simulations, and it represents the speed of the constant speed traveling wave.

### 4.1 An analytical argument for the stability of traveling wave speed

We introduced a small-parameter *ϵ* perturbation at t=0. This theoretical perturbation consisted of imposed initial conditions that force the wave to travel from x = −∞ to x= 0 at a constant speed faster than c_*fast*_. The initial wave speed is defined by

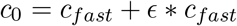

where a typical value for *ϵ* ≈ 0.01.

In this context, the consistency equation becomes:

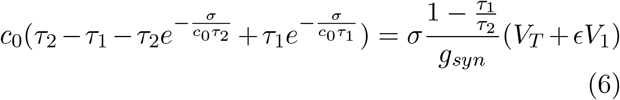

Figure 3 illustrates numerically how the perturbation imposed initial conditions that forced the wave to propagate from x= −∞ to x=0 at speed greater than c_*fast*_. The perturbation is removed at x=0, then the wave evolved due to the dynamics determined by the neuron and network parameters. The theoretical framework at hand depends on the magnitude of the small parameter *ϵ*. The accuracy of the evolution equations’ solutions decay as the magnitude of *ϵ* increases (Figure 4). Analysis of multiple simulations with increasing magnitudes of *ϵ* shows that the numerical results are in good agreement with the derivations’ solutions.

**Figure 3.**
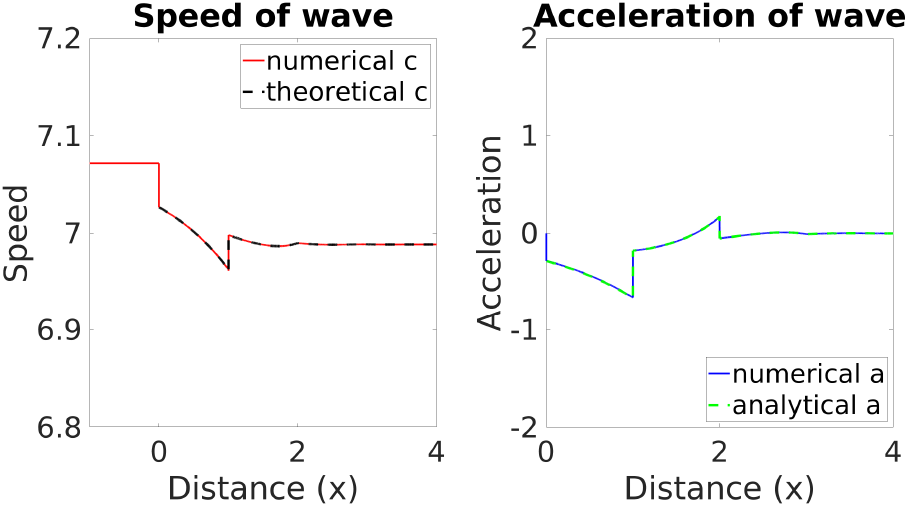
Faster wave speed perturbation: This graph shows the stability test of the wave speed solution c_*fast*_. Without loss of generality we consider a wave that prior to t=0 and x=0 travels at a slightly faster speed than the stable wave, that is, at *c*_0_ = *c_fast_* + *ϵ* ∗ *c_fast_*. The wave is initialized at a propagating speed faster than c_*fast*_, due to an external drive. The wave travels from *σ* = − inf to *σ*=0 at a constant speed *c*_0_ = *c_fast_* +*ϵ*∗*c_fast_*. At x=0, there is no external drive, and the wave is free to evolve due to the dynamics defined by network and neuron parameters. The wave shows damped oscillations while settling to c_*fast*_ propagating speed. The left panel displays traveling wave speed as a function of space. The red trace is the computer simulation, and the black trace is the result from Eq. 3 The right panel shows traveling wave acceleration as a function of space. The blue trace is the computer simulation, and the green trace is the solution from Eq 4. Note that acceleration is initially negative (the wave slows down), but then becomes positive because of the oscillations. Parameters: 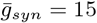, *σ* = 1, *τ*_1_ = 1, *τ*_2_ = 2, *V_T_* = 1, and *ϵ* = 0.012

**Figure 4.**
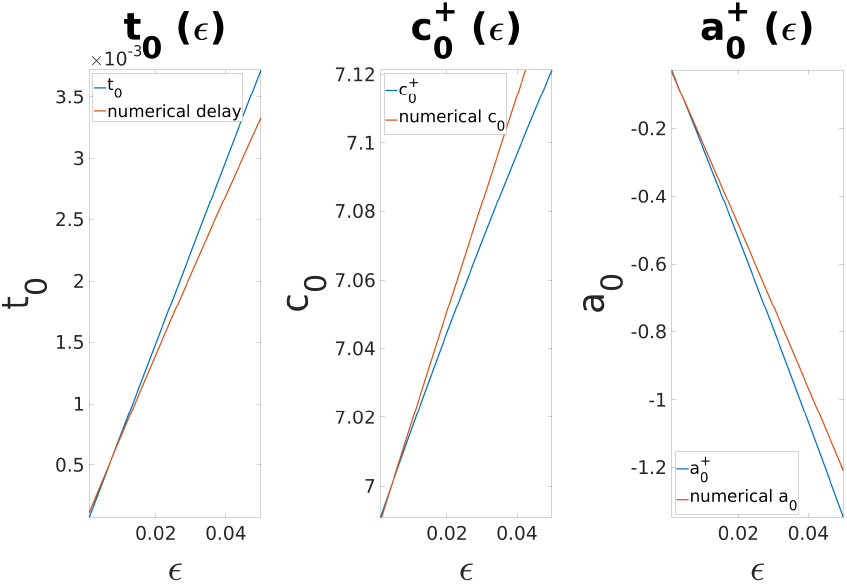
Correspondence between numerical results and analytical solutions. These graphs show the agreement between numerical results and the mathematical framework from the derivations. The left panel shows the delay computed from equation (7) and the delay from numerical simulations. The ratio of the slopes is 0.88 The middle panel shows the speed computed from equation (9) and the speed from numerical simulations. The ratio of the slopes is 0.84 The right panel shows the acceleration computed from equation (10) and the acceleration from numerical simulations. The ratio of the slopes is 0.89

At x=0 the perturbation is removed; here the traveling wave stopped and restarted propagation at a different speed after a delay. The delay is between the spiking time of the last neuron with the perturbation (x=0^−^), and the spiking time of the first neuron without perturbation (x=0^+^). The delay is a function of the voltage of the neuron in position x=0^+^, c_0_, the neuron, network parameters, and the firing times of neurons from x=−*σ* to x=0:

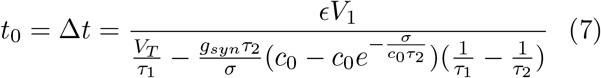

Our methods allowed us to estimate the wave characteristics: delay to initiate wave (t_0_), speed (c(0^+^)), and acceleration (a(0^+^)) at x=1.

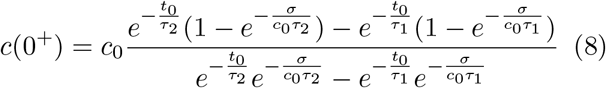

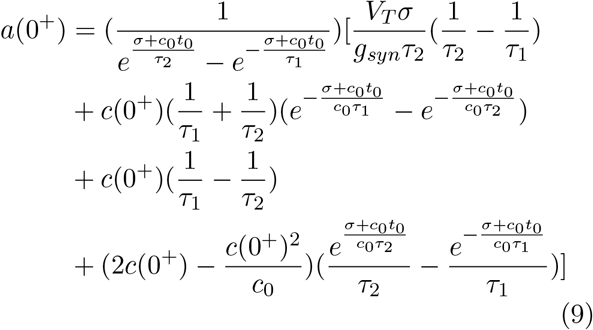

where

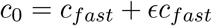

Figure 4 illustrates that the accuracy of our derivations depends on the magnitude of *ϵ*.

The slopes’ ratios demonstrate equations 7, 8, and 9 estimate the numerical delay, speed, and acceleration after the break with more than 80% accuracy. The discontinuity at the spatial boundary of the finite support connectivity kernel complicates solving analytically the evolution of the wave. In contrast to the exponential connectivity kernel where the absence of discontinuity within the connectivity kernel (one-way propagation) allows for explicit solutions [23].

The current framework assumes that propagation is monotonic from −∞ to +∞. Waves can accelerate or decelerate but not jump over any region. There is a delay for the fast wave case because the neuron at position *x* = 0^+^ needs time to integrate inputs before firing. But if we implement a slower wave perturbation (−*ϵ*), the neuron at position *x* = 0^+^ fires before the wave arrives and creates a second traveling wavefront. The equations and implementation of non-monotonic propagating waves with two wavefronts are beyond this paper’s scope.

### 4.2 Traveling wave stability: the effect of synaptic perturbations such as synaptic inhomogeneity, demyelination, and cell death

We investigate how perturbations to the microstructure of the neuronal network affect traveling wave propagation. In the synaptic inhomogeneity perturbation, the synaptic coupling parameter 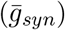 is allowed to oscillate sinusoidally as a function of space in a subsection of the network; the result is a sudden increase followed by a sharp decrease in synaptic conductance (Figure 5), which is one period of the sine function. In the demyelination perturbation, the synaptic coupling parameter decreases in a subsection of the network; here, the synaptic coupling is relatively weak (Figure 7). Lastly, in the cell death perturbation, the synaptic coupling is completely turned off for a subsection of the network; it emulates the network’s activity when the coupled neurons do not spike (Figure 9). If the subsection is not too large activity can jump over the dead region due to the longer-range connections that can extend over this region. We investigate how the traveling wave speed depends on the synaptic coupling parameter and space constant. Our results are numerical evidence of the stability of the traveling wave solution.

**Figure 5.**
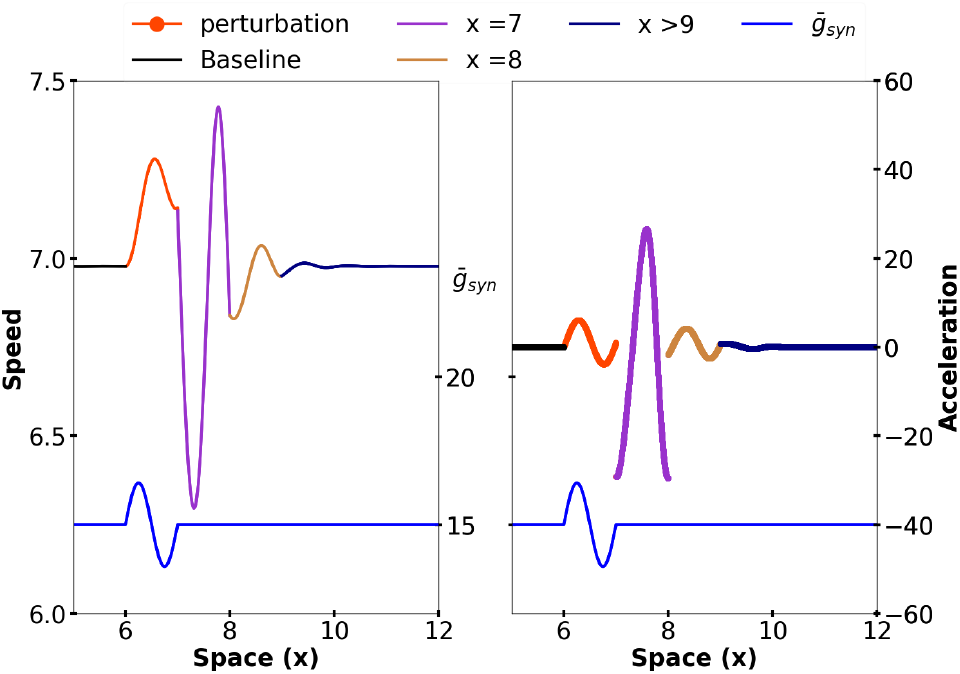
The wave speed oscillates as a function of both the synaptic coupling and the wave’s state. Both Panels: The wave travels from x= 0 to x=12. The blue line at the bottom of the graph represents how 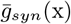 oscillates as a function of space and the sinusoidal perturbation, and is constant everywhere else. if x > 6 and x < 7

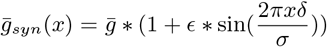

else

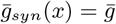 The traveling wave speed fluctuated dramatically while the synaptic inhomogeneity perturbation was present. Because of the effect of the delayed parameters of the finite support kernel, this perturbation produced a peak of traveling wave speed after the perturbation was removed. Interestingly, the magnitude of the second peak was greater than the first peak. The left panel illustrates that the speed of the wave at x =7 affects the speed of the wave at x=8; although the perturbation was totally removed at x>7. The wave speed peaks twice: The increase in local excitability 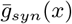 causes the first peak. The delayed effect of wave speed (*t*(*x*−*σ*), Eq. 3, *c*(*x*−*σ*), Eq. 4) causes the second peak. This phenomenon exemplifies some of the complicated dynamics of the finite support connectivity kernel compared to the exponential. The right panel illustrates how the traveling wave speed at any given “x” affects the instantaneous acceleration of the wave of subsequent “x+*σ*”. The instant acceleration fluctuated linearly relative to 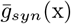. However, such covariation broke off after the removal of perturbation. Parameters: 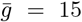, *σ* = 1, *τ*_1_ = 1, *τ*_2_ = 2, *V_T_* = 1, *δ* = 1*e^−^*^3^,and *ϵ* = 0.0943

**Figure 6.**
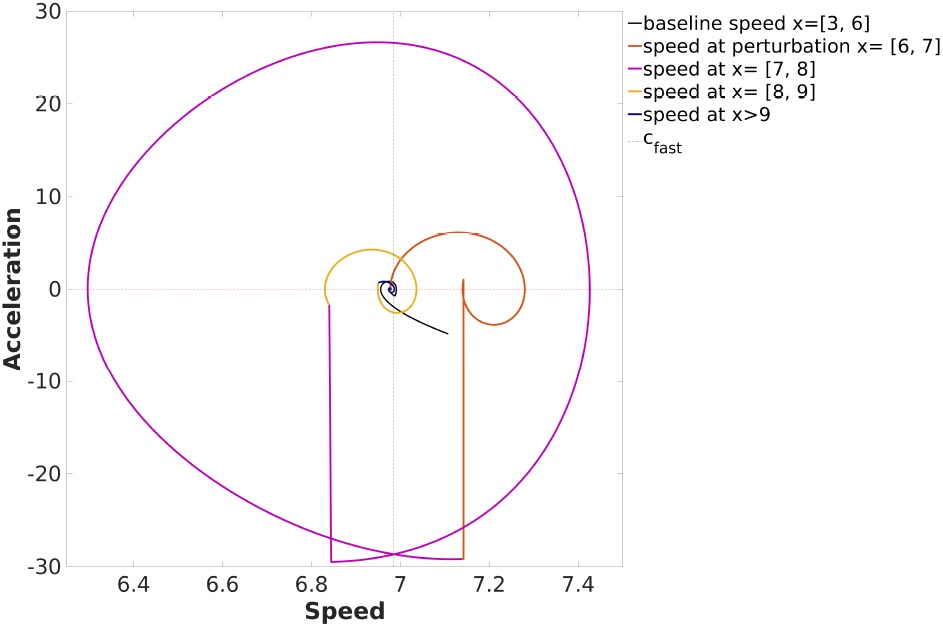
Co-evolution of traveling wave speed and acceleration in the presence of sinusoidal inhomogeneity perturbation. The traveling wave speed and acceleration evolve toward the intersection between c=c_*fast*_ and a=0. The black line represents the co-evolution of traveling wave speed and acceleration at baseline. The orange-red line represents the same variables but at the perturbation location. The violet, golden, and navy blue lines represent the wave’s evolution at 1, 2, and 3 or more synaptic space constants *σ* after removing the perturbation.

**Figure 7.**
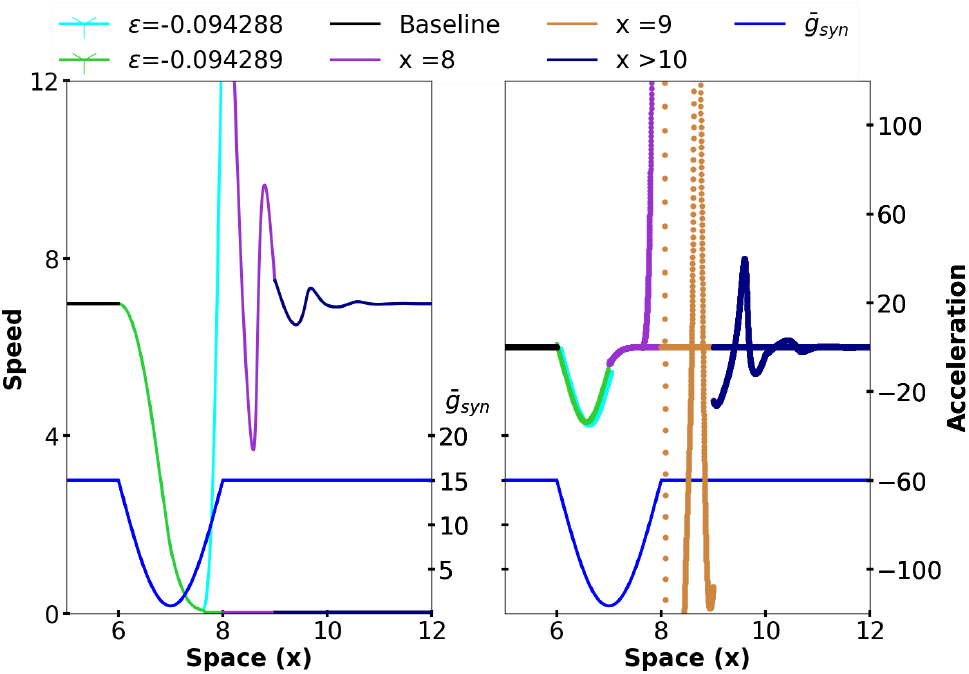
Neuronal traveling wave phenomena are all-or-none events: one small change in the control parameter separates activity propagation from propagation failure. Both Panels: The wave travels from x= 0 to x=12. The blue line at the bottom of the graph represents how 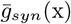 oscillates as a function of space and the demyelination perturbation. At the perturbation, 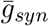 decreases sharply; the 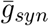 decay is determined by: if x > 6 and x < 7

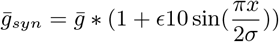

else

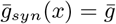 The cyan and purple wave speeds demonstrate how a small increase in parameter *ϵ* (+1e-6) determines whether the perturbation terminates the wave or if the wave continues to propagate after removing the perturbation. The amplitude of the perturbation controls the breaking point of the traveling wave. Here, two simulations with the same initial conditions and parameters but with one difference in *ϵ* (+1e-6) differ qualitatively. The cyan wave continued to propagate, at c_*fast*_ wave speed; the purple wave failed to propagate. Parameters: 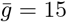, 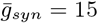, *σ* = 1, *τ*_1_ = 1, *τ*_2_ = 2, *V_T_* = 1, *ϵ*_1_ = −0.094289, *ϵ*_2_ = −0.094288.

**Figure 8.**
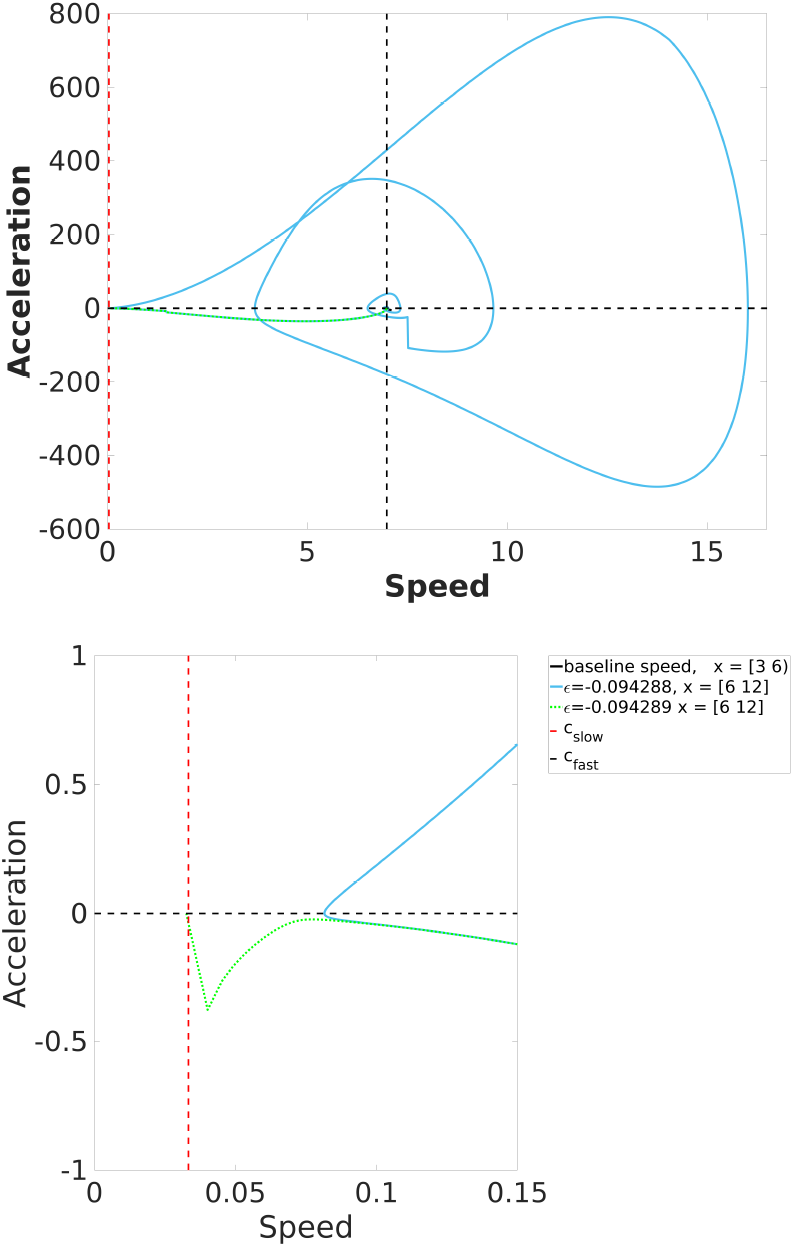
Co-evolution of traveling wave speed and acceleration in the presence of the demyelination perturbation. The left panel shows the evolution of traveling waves for two different values of *ϵ*: *ϵ* < *ϵ_critical_* (cyan), and *ϵ* > *ϵ_critical_* (green). The solid cyan line represents the co-evolution of the wave characteristics when the wave recovered successfully (*ϵ* < *ϵ_critical_*). The dotted green line represents the coevolution of traveling wave speed and acceleration when the wave failed to recover (*ϵ* > *ϵ_critical_*). The right panel illustrates the difference in the evolution of the green and cyan waves. We uniformly sampled one hundred points between *ϵ*=0 and −0.01; the critical value was between *ϵ*= −0.0942 and −0.0943. Again, we uniformly sampled one hundred points between *ϵ* =−0.0942 and −0.0943 to find *ϵ_critical_* with six-decimal point accuracy. For all *ϵ* < *ϵ_critical_*, the wave speed is never slower than *c_slow_*. However, for all *ϵ* > *ϵ_critical_*, the wave speed becomes slower than *c_slow_*.

**Figure 9.**
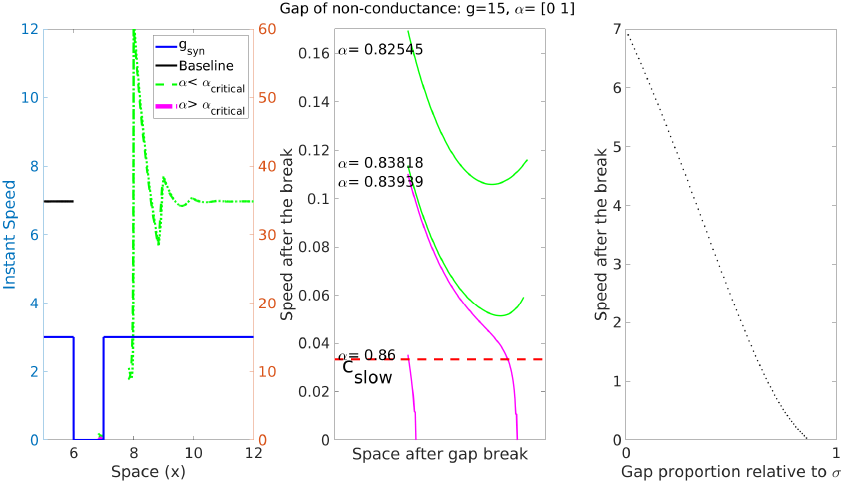
Dynamics of the traveling wave induced by a non-conducting gap. The simulation consists of a wave traveling at a constant speed *c_fast_*, located at x = 6 there is the non-conducting gap of “dead neurons”, which do not spike or synapse. The gap is relative to the synaptic space constant *σ*, and determined by the ratio: dead *gap* = *α* ∗ *σ* To compute *α*_*critical*_, we uniformly sampled one hundred points between 0 and 1. Then iterated between 0.8 and 0.86, to find *α*_*critical*_ with 4-point decimal accuracy. **Left Panel** The wave travels at a constant speed arriving at x = 6 (black trace). The green and magenta traces show the speed of the wave after the non-conducting gap results from multiple simulations; green curves represent *α* < *α*_*critical*_, and magenta represent *α* > *α*_*critical*_. As the non-conducting gap becomes larger, the wave speed after the break decreases. Larger gaps induce propagation failure. **Middle Panel** Speed after the gap for the simulations with *α* near *α*_*critical*_. The green and magenta traces show a critical qualitative change in traveling wave evolution as a result of a small parameter change. **Right Panel** The speed after the gap is a monotonically decreasing function of the gap size.

#### 4.2.1 Synaptic inhomogeneity perturbation

Synaptic inhomogeneity refers to the media microstructure; it represents periodic variation of strenghts of synaptic coupling. We assume 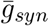 is equal throughout the network, except for one synaptic space constant (*σ*), where 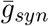 oscillates sinusoidally with wavelength *λ* = *σ*. Here, the connections between neurons are stronger, followed by weaker connections than baseline (Figure 5). These synaptic coupling patterns are common in neuronal networks where neurons from different populations co-exist, such as optical preference columns in the visual cortex and the barrel cortex [25]. In our simulations, these synaptic arrangements show interesting transient phenomena while also supporting the stability of the traveling wave speed c_*fast*_ (Figure 6). The traveling wave propagated at a constant speed before the perturbation. The wave speed oscillated at the perturbation location; the speed and acceleration drifted away from the stable baseline state. After removing the perturbation, the wave speed continued to oscillate. The oscillations damped while approaching c_*fast*_ (Figure 5); while the wave speed approached c_*fast*_, the wave acceleration approached zero. The wave reaches a constant speed when speed equals c_*fast*_ and acceleration is 0. Interestingly, the effects of the perturbation are maximal during the next two synaptic footprints to the right of the perturbation region.

#### 4.2.2 Demyelination perturbation

Synaptic demyelination is a neurodegenerative condition in which neurons lose their myelin sheath. Demyelination may be caused by aging [26], Huntington’s disease [27], and ALS [28]. To model it, we assume 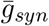 is equal throughout the network, except for two synaptic space constants, where 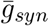 oscillates quadratically with wavelength *λ* = *σ* (Figure 7). In the demyelination perturbation, the strength of the neuronal connections decreases dramatically in a network subsection. The decrease in 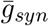 synaptic coupling represents the decay of synaptic coupling is common in demyelinating and neurodegenerative diseases [26–28]. We performed simulations to investigate the traveling wave speed and acceleration with varying magnitudes of perturbations. Our results strongly suggest that for this class of models, neuronal traveling wave phenomena are all-or-none events (Figure 7).

The traveling wave propagated at a constant speed before reaching the location of the perturbation. At the perturbation location, 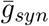 decays drastically for one *σ*; in the subsequent *σ*, 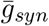 rapidly recovers baseline (Figure 7). The wave traveled at a constant speed when it arrived at the perturbation location. Our simulations showed a critical value of *ϵ* (Figure 8) that separated the waves that recovered from those that died off. The waves that recovered after removing the perturbation oscillated while approaching c_*fast*_; however, the waves that died off approached c_*slow*_ (Figure 8). Similar to other models, propagation failure occurs suddenly as the parameter *ϵ* decreases a certain threshold. In contrast to other models, such as the exponential kernel, propagation failure occurs quickly instead of at farther away distances.

#### 4.2.3 Cell-death perturbation

We assume synaptic coupling is equal throughout the network, except for a subsection where neurons are not coupled and do not spike. As the wave travels from −∞ toward positive values for ‘x’ it encounters a non-conducting gap for a finite region, then synaptic coupling returns to baseline values. The gap’s size is determined by the ratio *α* ∗ *σ*, where the value of *α* ranges from 0 to 1. In the cell-death perturbation, a gap of synaptic non-conductance represents the area of dead neurons. The perturbation methods model acute events of neural cell death that may result from cerebral infarction [29] or traumatic brain injury [30,31]; these sorts of insults produce neuronal cell death at the location of the accident, while adjacent neurons may survive the insult.

Similar to the other section results, the outcomes from the cell-death perturbation support the idea that constant speed neural waves are all-or-none events (Figure 9). Waves typically recover from smaller non-conducting gaps; the wave speed oscillates transiently but eventually settles back to a constant speed value, namely c_*fast*_ (*α* < *α*_*critical*_, Figure 9 green). However, larger perturbations break traveling wave propagation (*α* >= *α*_*critical*_, Figure 9 magenta). The traveling wave’s speed is slower than c_*fast*_ right where synaptic coupling returns to baseline; the speed after the break decreases monotonically as *α* grows and approaches *α*_*critical*_.

Computer simulations determined the smallest *α*_*critical*_ ∗ *σ* gap that breaks traveling wave propagation (Figure 10). *α*_*critical*_ represents the ratio of the synaptic space constant *σ* in the borderline between traveling wave propagation and failure. Simulations with varying values of 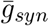 demonstrate that *α*_*critical*_ is relatively low at low synaptic conductances, but *α*_*critical*_ increases rapidly, and approaches asymptotically to *σ* as synaptic coupling grows. Indicating that as excitation in the network increases propagation failure is less likely to be triggered, as it requires larger perturbations to suppress the activity in some propagation regions. Figure 10 shows a perfect fit between 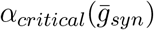 and the power function. Interestingly, this is close to a fit of the form 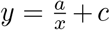, because the exponent is close to −1. There is an inverse relationship between 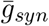 and *α*_*critical*_.

**Figure 10.**
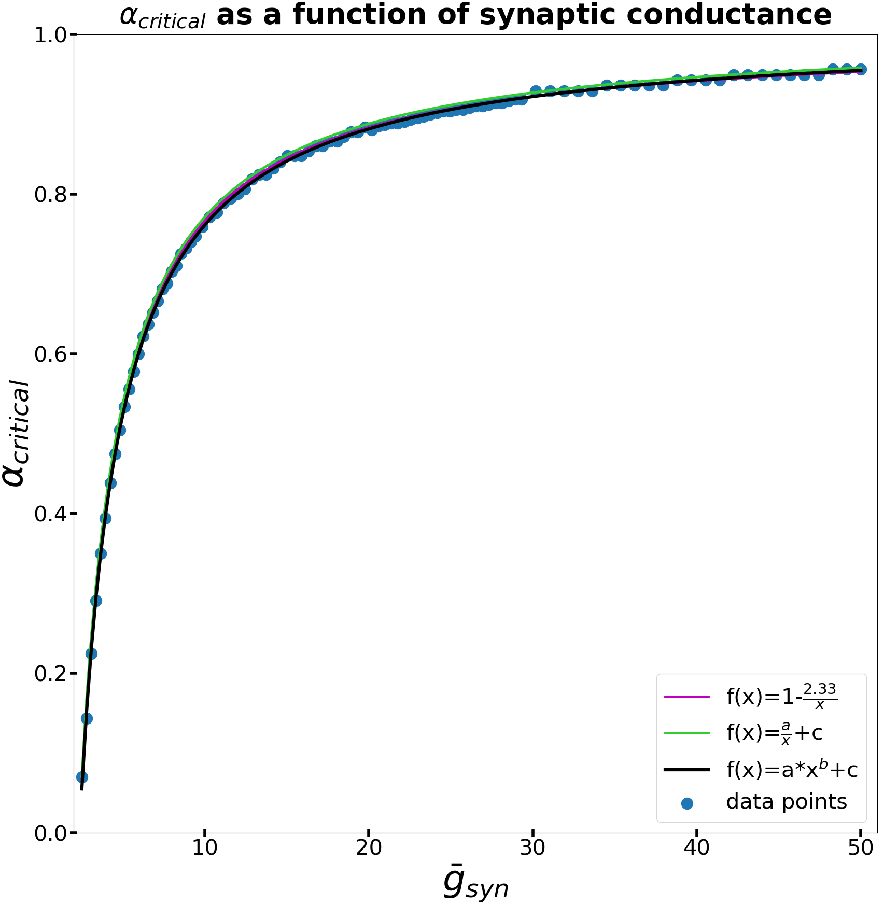
The smallest gap lengths that induce propagation failure gap=*σα*. Networks with strong excitatory coupling allow for robust wave propagation. The curve above shows as *g*_*syn*_ increases, *α*_*critical*_ must also increase to produce propagation failure. Intuitively, this indicates that when overall network excitation is larger, more drastic reduction in the non-conducting gap is needed to produce propagation failure. Curve fit analysis demonstrated the power function was a good fit for the curve.

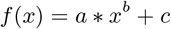 Coefficients (with 95% confidence bounds): a = −2.336 (−2.357, −2.315) b = −0.982 (−0.9904, −0.9736) c = 1.005 (1.003, 1.007) SSE = 0.0008004, *R*^2^ = 0.9998, Adjusted R^2^ = 0.9998, RMSE = 0.002918 Interestingly, the exponent is so close to −1 that other curves such as

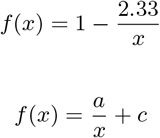

may also fit the asymptotic trend.

## 5 Conclusions and future directions

We present an extensive analysis of traveling waves in neuronal networks with finite support synaptic coupling. In neuronal networks with exponential or Gaussian connectivity kernels [23], neurons are connected to neighbors up to infinity; although the connections decay with distance. In contrast, these model networks represent a simplification in network topology by assuming that synapses strictly space-dependent. Connections are only between neighboring neurons within a synaptic footprint *σ*, and zero elsewhere; this simplification is based on the assumption that synaptic coupling is strictly space-dependent. However, the finite support kernel introduces discontinuities in the network. The finite support neuronal network proposes a more straightforward mathematical function and biological structure compared to other connectivity functions, such as Gaussian or exponential. Despite the inherent simplicity in the finite support function, the mathematical model is complicated and has no analytical solutions. However, we managed to describe wave propagation as a set of evolution equations that predict wave dynamics without the need to run the entire computer simulation equations (2) (3) and (4). In order to investigate the stability of the wave, we presented a number of perturbations. First, a traveling wave coming from −∞ to 0 at a constant speed *c*_0_ = *c_fast_* +*ϵc_fast_*. Simulations and analytical studies show there is a delay to spike once the perturbation is removed. We compute the delay, the speed, and acceleration after the break, equations (7), (8), and (9) respectively. We then presented perturbations relevant to a biological context, resembling effects of demyelinating disorders and cell death. Demyelinating disorders were presented as continuous perturbations (Figure 5, and Figure 7) in the topology of parameter *g*_*syn*_ in a subsection the network. These results were consistent with the notion that synaptic coupling facilitates traveling wave propagation: in the cases the introduced perturbation increased coupling, the wave accelerated; and in the cases the perturbation decreased coupling, the wave decelerated. If the wave decelerates slower than *c_slow_*, propagation fails to continue (Figure 7). The last perturbation we presented was a non-conducting gap, which resembles a small section of “dead tissue” (Figure 9). The gap was defined as *gap* = *ασ* and represented the ratio *gap* = *α* ∗ *σ*. For any given *g*_*syn*_ there is a corresponding critical value *ϵ_critical_* between activity propagation and failure (Figure 10). The demyelination and dead tissue perturbations (Figures 7 and 9, respectively) show remarkable stability of the system because despite how much the perturbation decelerated the wave, all the waves that continued to propagate evolved back to the steady-state *c_fast_*.

Altogether, the present work demonstrates waves respond robustly in neuronal networks with finite support coupling and that network parameters influence traveling wave propagating speed. Waves in neuroscience are ubiquitous, and unraveling population dynamics can inform us of underlying mechanisms that give rise to neuronal function and dysfunction. For example, the smooth perturbation (Figure 7) represents decreasing synaptic coupling in a network subpopulation. This phenomenon is common in myelination diseases, such as Huntington’s Disease and multiple sclerosis. In this framework, it is intuitive to understand why many symptoms are not expressed until demyelination reaches a critical point, at which neuronal function fall apart. Likewise, in the non-conducting gap of “dead tissue” (Figure 9) may be an example of post-infarction neuronal tissue. This hypothesis could explain the impairment of neuronal functions in context of the local characteristics and topology of the network where the cerebral infarction took place. The exponential connectivity kernel imposes the longest-reaching connections (*e*^−*x*^) [23]. The finite support kernel imposes the shortest ranging connections. Future studies should focus on the Gaussian kernel, which imposes midrange connections 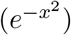. However, the derivation of the evolution equations become more complicated for the Gaussian kernel inside the integral. The second derivative of the Gaussian kernel results in a polynomial times the Gaussian. As a result, the system of equations becomes harder to solve.

## References

[1] T. K. Sato, I. Nauhaus, and M. Carandini, “Traveling Waves in Visual Cortex,” Neuron, vol. 75, pp. 218–229, Jul 2012.

[2] T. Wanger, K. T., M. T. Lippert, J. Goldschmidt, and F. W. Ohl, “Wave propagation of cortical population activity under urethane anesthesia is state dependent,” BMC Neuroscience, vol. 14, p. 78, Dec 2013.

[3] J. B. Ackman and M. C. Crair, “Role of emergent neural activity in visual map development,” Curr. Opin. Neurobiol., vol. 24, pp. 166–175, Feb 2014.

[4] J. B. Ackman, T. J. Burbridge, and M. C. Crair, “Retinal waves coordinate patterned activity throughout the developing visual system,” Nature, vol. 490, p. 219, Oct 2012.

[5] T. P. Zanos, P. J. Mineault, K. T. Nasiotis, D. Guitton, and C. C. Pack, “A Sensorimotor Role for Traveling Waves in Primate Visual Cortex,” Neuron, vol. 85, pp. 615–627, Feb 2015.

[6] I. Nauhaus, L. Busse, D. L. Ringach, and M. Carandini, “Robustness of Traveling Waves in Ongoing Activity of Visual Cortex,” J. Neurosci., vol. 32, pp. 3088–3094, Feb 2012.

[7] A. Compte, M. V. Sanchez-Vives, D. A. McCormick, and X.-J. Wang, “Cellular and network mechanisms of slow oscillatory activity (<1 Hz) and wave propagations in a cortical network model.,” undefined, 2003.

[8] M. Murakami, H. Kashiwadani, Y. Kirino, and K. Mori, “State-Dependent Sensory Gating in Olfactory Cortex,” Neuron, vol. 46, pp. 285–296, Apr 2005.

[9] A. Reimer, P. Hubka, A. K. Engel, and A. Kral, “Fast Propagating Waves within the Rodent Auditory Cortex,” Cereb. Cortex, vol. 21, pp. 166–177, Jan 2011.

[10] M. Chrostowski, L. Yang, H. R. Wilson, I. C. Bruce, and S. Becker, “Can homeostatic plasticity in deafferented primary auditory cortex lead to travelling waves of excitation?,” J. Comput. Neurosci., vol. 30, pp. 279–299, Apr 2011.

[11] D. R. Belov, P. A. Stepanova, and S. F. Kolodyazhnyi, “Traveling Waves in the Human EEG during Voluntary Hand Movements,” Neurosci. Behav. Physi., vol. 45, pp. 1043–1054, Nov 2015.

[12] D. Rubino, K. A. Robbins, and N. G. Hatsopoulos, “Propagating waves mediate information transfer in the motor cortex,” Nat. Neurosci., vol. 9, p. 1549, Nov 2006.

[13] S. Dipierro and E. Valdinoci, “A Simple Mathematical Model Inspired by the Purkinje Cells: From Delayed Travelling Waves to Fractional Diffusion,” Bull. Math. Biol., vol. 80, pp. 1849–1870, Jul 2018.

[14] D. B. Poll and Z. P. Kilpatrick, “Velocity Integration in a Multilayer Neural Field Model of Spatial Working Memory,” SIAM J. Appl. Dyn. Syst., Jul 2017.

[15] K. Takahashi, M. Saleh, R. D. Penn, and N. Hatsopoulos, “Propagating Waves in Human Motor Cortex,” Front. Hum. Neurosci., vol. 5, Apr 2011.

[16] A. Bahramisharif, M. A. J. van Gerven, E. J. Aarnoutse, M. R. Mercier, T. H. Schwartz, J. J. Foxe, N. F. Ramsey, and O. Jensen, “Propagating Neocortical Gamma Bursts Are Coordinated by Traveling Alpha Waves,” J. Neurosci., vol. 33, pp. 18849–18854, Nov 2013.

[17] M. Massimini, R. Huber, F. Ferrarelli, S. Hill, and G. Tononi, “The Sleep Slow Oscillation as a Traveling Wave,” J. Neurosci., vol. 24, pp. 6862–6870, Aug 2004.

[18] E. V. Lubenov and A. G. Siapas, “Hippocampal theta oscillations are travelling waves,” Nature, vol. 459, p. 534, May 2009.

[19] H. Zhang and J. Jacobs, “Traveling Theta Waves in the Human Hippocampus,” J. Neurosci., vol. 35, pp. 12477–12487, Sep 2015.

[20] C. A. Cuellar, J. A. Tapia, V. Juárez, J. Quevedo, P. Linares, L. Martínez, and E. Manjarrez, “Propagation of Sinusoidal Electrical Waves along the Spinal Cord during a Fictive Motor Task,” J. Neurosci., vol. 29, pp. 798–810, Jan 2009.

[21] A. Laan, T. Gutnick, M. J. Kuba, and G. Laurent, “Behavioral Analysis of Cuttlefish Traveling Waves and Its Implications for Neural Control,” Curr. Biol., vol. 24, pp. 1737–1742, Aug 2014.

[22] D. Kleinfeld, K. R. Delaney, M. S. Fee, J. A. Flores, D. W. Tank, and A. Gelperin, “Dynamics of propagating waves in the olfactory network of a terrestrial mollusk: an electrical and optical study,” Journal of Neurophysiology, Sep 1994.

[23] J. Zhang and R. Osan, “Analytically tractable studies of traveling waves of activity in integrate-and-fire neural networks,” Phys. Rev. E, vol. 93, p. 052228, May 2016.

[24] R. Osan and B. Ermentrout, “The evolution of synaptically generated waves in one- and two-dimensional domains,” Physica D, vol. 163, pp. 217–235, Mar 2002.

[25] E. R. Kandel, J. H. Schwartz, T. M. Jessell, D. of Biochemistry, M. B. T. Jessell, S. Siegelbaum, and A. Hudspeth, Principles of neural science, vol. 4. McGraw-hill New York, 2000.

[26] A. M. Adinolfi, J. Yamuy, F. R. Morales, and M. H. Chase, “Segmental demyelination in peripheral nerves of old cats,” Neurobiology of aging, vol. 12, no. 2, pp. 175–179, 1991.

[27] O. Phillips, C. Sanchez-Castaneda, F. Elifani, V. Maglione, A. Di Pardo, C. Caltagirone, F. Squitieri, U. Sabatini, and M. Di Paola, “Tractography of the corpus callosum in huntingtons disease,” PloS one, vol. 8, no. 9, p. e73280, 2013.

[28] S. H. Kang, Y. Li, M. Fukaya, I. Lorenzini, D. W. Cleveland, L. W. Ostrow, J. D. Rothstein, and D. E. Bergles, “Degeneration and impaired regeneration of gray matter oligodendrocytes in amyotrophic lateral sclerosis,” Nature neuroscience, vol. 16, no. 5, pp. 571–579, 2013.

[29] I. Ferrer and A. M. Planas, “Signaling of cell death and cell survival following focal cerebral ischemia: life and death struggle in the penumbra,” Journal of Neuropathology & Experimental Neurology, vol. 62, no. 4, pp. 329–339, 2003.

[30] R. Raghupathi, “Cell death mechanisms following traumatic brain injury,” Brain pathology, vol. 14, no. 2, pp. 215–222, 2004.

[31] A. Rink, K.-M. Fung, J. Q. Trojanowski, V. Lee, E. Neugebauer, and T. K. McIntosh, “Evidence of apoptotic cell death after experimental traumatic brain injury in the rat.,” The American journal of pathology, vol. 147, no. 6, p. 1575, 1995.

